# MuPET-Flow: Multiple Ploidy Estimation Tool from Flow cytometry data

**DOI:** 10.1101/2024.01.24.577056

**Authors:** C. Gómez-Muñoz, G. Fischer

## Abstract

**Summary:** Ploidy, representing the number of homologous chromosome sets, can be estimated from flow cytometry data acquired on cells stained with a fluorescent DNA dye. This estimation relies on a combination of tools that often require scripting, individual sample curation, and additional analyses. To automate the ploidy estimation for multiple flow cytometry files, we developed MuPET-Flow—a Shiny graphical user interface tool. MuPET-Flow allows users to visualize cell fluorescence histograms, detect the peaks corresponding to the different cell cycle phases, perform a linear regression using standards, make ploidy or genome size predictions, and export results as figures and table files. The tool was benchmarked with known ploidy datasets, yielding consistent ploidy results. MuPET-Flow produces ploidy results comparable to those from other available tools; however, MuPET-Flow stands out for its user-friendly interface, which allows the whole process to be carried out in one place, considerably speeding up analysis, particularly in projects involving large numbers of samples.

**Availability and implementation:** https://github.com/CintiaG/MuPET-Flow.

## INTRODUCTION

Ploidy (n) is the number of homologous chromosome sets in a cell. Cells can be haploid if they have one set (1n), diploid if they have two (2n), or polyploid if they have three or more (≥3n). Polyploidy exists in several branches of the tree of life (Otto and Whitton, 2000), notably in the *Saccharomyces* genus (Peter *et al*., 2018; Gómez-Muñoz *et al*., 2021), and in plants (Čertner *et al*., 2022).

Flow cytometry (FC) allows measuring the fluorescence of single cells stained with a DNA intercalant, such as propidium iodide (PI). When this fluorescence is plotted against the cell count (histogram of fluorescence intensity), usually two peaks corresponding to the different cell cycle stages, G0/G1 and G2, are recognizable. The peaks’ fluorescence intensity is proportional to the DNA content, which allows inferring both ploidy and genome size. To achieve this, the peaks’ fluorescence of query samples needs to be identified and then compared to that of known standards. Such standards can be run separately (external standardization) or within the sample (internal standardization) (Todd *et al*., 2018; Sliwinska *et al*., 2022).

Several software tools for FC data analysis exist. Proprietary software, such as FlowJo (Becton Dickinson & Company, USA), are usually tied to the equipment or require payment. Open-source software, developed in R and Python programming languages, are freely available and have the potential to automate the production of publication-ready images but typically lack a graphical user interface (GUI) and involve scripting, which might be daunting for some users. For the specific case of peaks detection and ploidy estimation, two open-source tools deserve special mention as they offer a GUI. On one hand, Cytoflow (Teague, 2022), implemented in Python, is a versatile FC data analysis package. In this software, data can be loaded and visualized through a GUI. However, it requires manual configuration, and the formal analysis must be conducted outside the software (Table S1). On the other hand, flowPloidy, implemented in R (Smith *et al*., 2018), is a tool that provides powerful peaks and debris fitting, as well as gating, facilitated by the inclusion of an R Shiny GUI (Chang *et al*., 2023). Nevertheless, it still involves scripting, and individual sample curation (Table S1).

Given the importance of FC in ploidy estimation and the limitations of the existing tools, we developed MuPET-Flow (Multiple Ploidy Estimation Tool from Flow cytometry data). MuPET-Flow is a GUI tool that automates file uploading and configuration, peak fluorescence intensity detection, multiple histogram visualizations, peak error curation, ploidy and genome size calculations, and easy results export (Table S1).

We benchmarked our tool using *Saccharomyces* datasets and demonstrated its applicability in plant species for ploidy and genome size estimation. Additionally, we compared MuPET-Flow to Cytoflow and flowPloidy. MuPET-Flow shares several features and advantages with these existing tools (Table S1), but its novelty consists in automating and combining features that usually exist separately, as discussed below. Notably, MuPET-Flow is the only tool that allows full and simple in-app calculation of ploidy. MuPET-FC is available at https://github.com/CintiaG/MuPET-FC.

## MATERIALS AND METHODS

### Flow cytometry data acquisition

*S. cerevisiae* strains included two natural polyploids CLQCA_17-111 (abbreviated as CRE; 3n) and CH10 (abbreviated as AVQ; 5n), used as test, and four laboratory strains, BY4742 (1n), BY4743 (2n), YPS128_3n (3n), and YPS128_4n (4n), serving as standards. Three biological replicates of yeast cells were cultured in YPD media (Sigma-Aldrich, USA), fixed in ethanol 70% and washed twice with Phosphate-buffered saline solution 1X. Subsequently, the cells were treated with RNAse A (EUROMEDEX, France) at 1 mg/mL at 37°C for 2h, and stained with PI (Thermo Fisher Scientific, USA) at 50 µg/mL. Fluorescence was measured in the MACSQuant VYB system (Miltenyi Biotec, Germany), in channel FL4-A (605-625 nm). Files were gated by FSC-A vs SSC-A and FSC-A vs FSC-H using CytoExploreR (Hammill, 2021). The FC files generated are available on MuPET-Flow’s GitHub. *Saccharomyces pastorianus* files were obtained from Gómez-Muñoz *et al*. (2021), while *Solanum pseudocapsicum* files (Čertner *et al*., 2022) were downloaded from the FlowRepository (ID FR-FCM-Z45W; Spidlen *et al*., 2012).

### Tool description and performance evaluation

MuPET-Flow is implemented in the R package Shiny (Chang *et al*., 2023) and comprises three tabs (Fig. 1). On the first tab, ‘Peaks’, multiple local Flow Cytometry Standard (FCS) files can be uploaded through the GUI using internally flowCore (Hahne *et al*., 2009). MuPET-Flow automatically scans the files for available channels and displays them in and dropdown menu, from which one must be selected. The tool then estimates the number of breaks (minimum 256) and automatically generates a histogram in the chosen channel for all samples. A ‘Local Polynomial Regression Fitting’ is applied for histograms smoothing (default 0.5). Peaks are detected using a local maximum algorithm by employing a sliding window of a given width (default 50). Optionally, individual files can be selected through a dropdown menu to display the sample’s histogram and peak detection results, and the smoothing level and window width adjusted by a numeric input menu. The minimum cell count to call peak can be selected (default 5). The algorithm assumes that only the G0/G1 and G2 populations exist, and only two peaks can be used for the subsequent steps. However, additional detected peaks can be explored and selected through a checkbox.

**Fig. 1.**
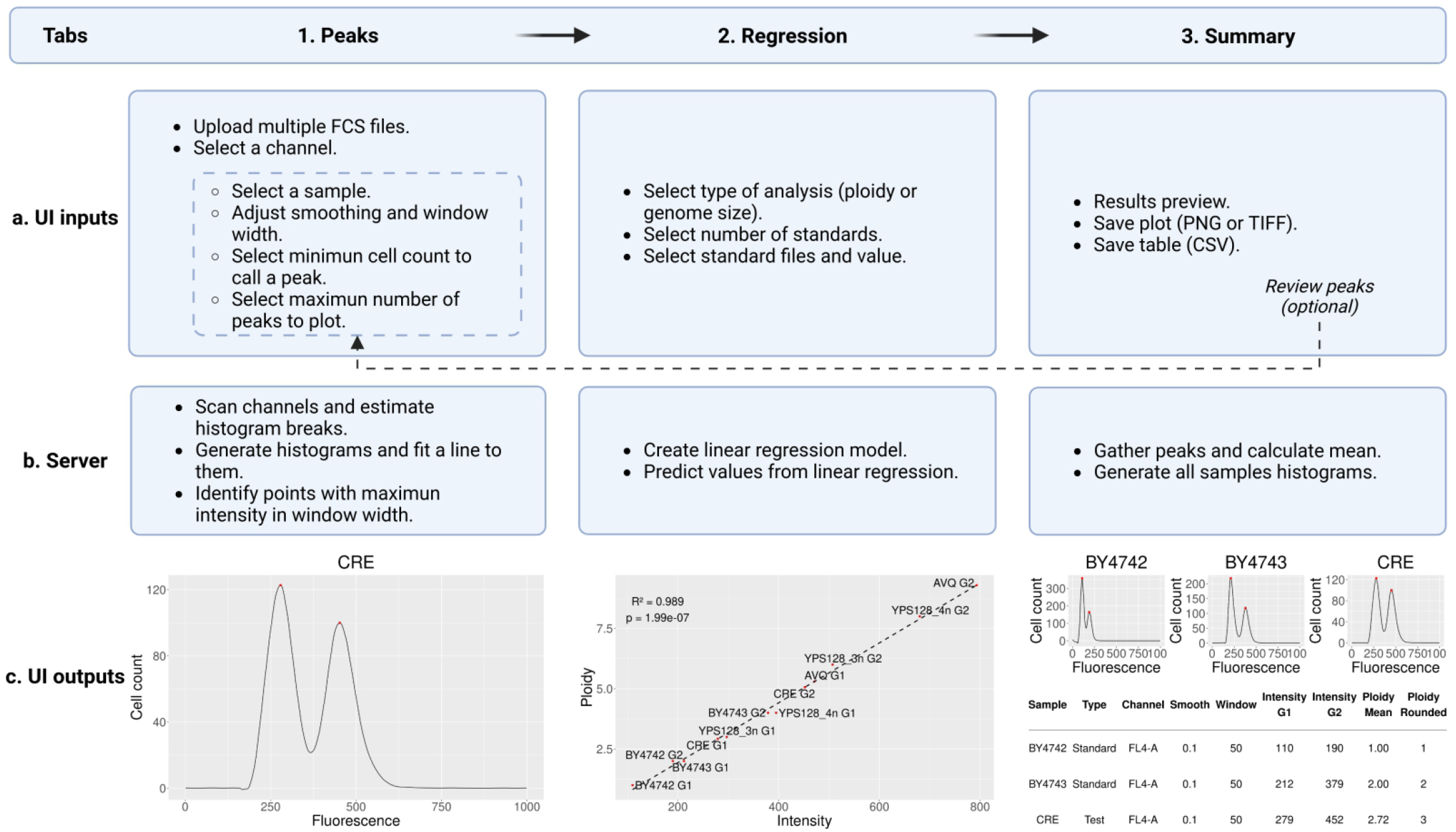
MuPET-Flow: A Shiny app for estimating ploidy in multiple samples. The MuPET-Flow workflow is divided into three tabs: ‘Peaks’, ‘Regression’, and ‘Summary’. The main functionalities consist of User Interface (UI) inputs, processes executed in the Server, and UI outputs. Adjusting the inputs in the dashed box is optional. Created with BioRender.com.

In the second tab, ‘Regression’, MuPET-Flow estimates ploidy or genome size using a linear regression based on external standards. Therefore, the type of analysis, the number of standards files, and their corresponding values should be specified. A minimum of two distinct standards is necessary. The two peaks of the standards are considered for linear regression, and the ploidy or genome size of both peaks of the test samples is predicted. The linear regression results are displayed in a text box.

Finally, all generated histograms can be previewed in the third tab called ‘Summary’. In case not all of the samples were individually inspected and an error is found, it is possible to return to the first tab to review any parameters and selected peaks, provided that regression is redone as well. All used parameters, such as ‘smoothing’ and ‘window’, as well as the mean ploidy or genome size (presented as decimal and rounded values), are reported. The histograms can be saved as a figure (PNG or TIFF), and the table as a comma-separated value file (CSV).

The peaks’ fluorescence of the *S. cerevisiea* dataset was also determined using Cytoflow (Teague, 2022) and flowPloidy (Smith *et al*., 2018), following their instructions. The ploidies were obtained by applying the same principle as MuPET-Flow (linear regression). These results were compared with a paired Wilcoxon signed rank exact test. To assess the tool’s execution time, we generated a mock dataset by duplicating the files of the six *S. cerevisiae* analyzed strains for all replicates resulting in 36 FCS files (considered a manageable number for visualization) with a total size of 119 MB. Using the R ‘system.time’ function, the user, system, and elapsed mean times of three runs were obtained for the main application processes. Full computer specifications for developing and testing MuPET-Flow are in Table S2.

## RESULTS AND DISCUSSION

### Benchmarking with newly generated *S. cerevisiae* FC data

To test our application, we determined the ploidy of two *S. cerevisiae* polyploids, CRE and AVQ, from newly obtained PI-stained cells FC data. The histograms of the two tested strains and four standards were visualized simultaneously (Fig. S1), and the smoothing and window were adjusted for a few samples (Table S3). The estimated peaks’ intensities and the inferred ploidy by linear regression are reported in Table S3. The fluorescence intensity varied among the biological replicates; however, there was a good agreement in the mean ploidy for CRE and AVQ in the triplicates. Similarly, the rounded ploidy of the tested strains was identical to that reported by Peter *et al*. (2018). Additionally, we examined the execution time over the *S. cerevisiea* mock dataset consisting of 36 files with MuPET-Flow. The total user time for MuPET-Flow’s main processes was less than 2 seconds (Table S2).

### App testing with published data in other species

To further assess the app’s performance, we challenged it with FC data originating from different instruments, fluorophore technology, and species. We utilized two previously published cytometry datasets, one from the yeast *S. pastorianus* (Gómez-Muñoz *et al*., 2021) for genome size estimation and another from the plant *S. pseudocapsicum* (Čertner *et al*., 2022) for ploidy determination. In the *S. pastorianus* dataset, we obtained a genome size of the query strain *S. pastorianus* 790 of 58.64 Mb, close to the 60.1 Mb previously estimated (Table S4; Gómez-Muñoz *et al*., 2021). For *S. pseudocapsicum*, multiple peaks were detected in the fruit skin file. The ploidy of the highest intensity peaks was investigated, revealing correspondence with the 8n peak previously reported for this tissue (Table S5; Čertner *et al*., 2022). Thus, MuPET-Flow is primarily developed to estimate ploidy in yeast; however, it can also be applied to measure genome size and in other species.

### MuPET-Flow comparison with open-source tools for ploidy estimation

To emphasize the advantages of MuPET-Flow’s pipeline, we compared it with the pipelines of Cytoflow and flowPloidy, focusing on their ability to calculate ploidy over the *S. cerevisiea* dataset. We examined whether the peaks’ fluorescences and ploidies obtained with MuPET-Flow differed from those obtained with Cytoflow and flowPloidy. We observed a significant difference (p < 0.05) in the peaks’ fluorescence between MuPET-Flow and flowPloidy, as well as between Cytoflow and flowPloidy, but not between MuPET-Flow and Cytoflow. However, no differences were observed in the inferred ploidy values across the three tools (Table S6).

Compared to MuPET-Flow, Cytoflow (Teague, 2022) requires extensive manual configuration and using the 1D mixture model for detection of the different peaks’ fluorescence. However, the number of peaks for the model needs to be specified, and the same number is applied to all samples. Additionally, for ploidy estimation, the data needs to be exported and analyzed outside the software (Table S1). Likewise, FlowPloidy explicitly necessitates detecting channels, specifying the number of breaks for histograms, and saving the images via scripting. Furthermore, failed models needed to be inspected and corrected individually. Finally, flowPloidy is principally intended for calculating plant genome size with internal standards. When external standards are used, the peaks’ means need to be exported and the ploidy calculated through additional operations (Table S1).

On the contrary, MuPET-Flow is an easy-to-use tool that assembles and automates all necessary actions for ploidy estimation (Table S1). For instance, MuPET-Flow allows the visualization of samples both individually and simultaneously, as in Cytoflow, and the capability to correct histograms, somewhat similar to flowPloidy. These characteristics combined facilitates the identification and curation of the few problematic samples, not feasible in Cytoflow, and more efficient than flowPloidy which necessitates inspecting every one of them. Furthermore, MuPET-Flow obtained histograms and peaks’ intensities can be saved as a figure and a table, always with the aid of a GUI (Table S1). An additional feature of MuPET-Flow is the ability to detect multiple peaks due to its local maxima algorithm, without prior knowledge of the number of peaks. The multiple peaks present in certain samples can be explored and selected, as observed in the case of endopolyploidy reported in *S. pseudocapsicum* (Čertner *et al*., 2022).

Cytoflow, flowPloidy, and MuPET-Flow individually present several advantages and disadvantages (Table S1). Nevertheless, MuPET-Flow is an attractive option as it automates and facilitates many steps in ploidy estimation. It is the only tool capable of performing this process within the app in a straightforward manner, while reducing the need for manual curation. Overall, MuPET-Flow minimizes the time required for ploidy analysis, which is particularly beneficial for projects dealing with a large number of samples routinely.

## WEB RESOURCES

MuPET-FC code and execution instructions are available at https://github.com/CintiaG/MuPET-FC.

## DATA AVAILABILITY STATEMENT

*S. cerevisiae* dataset is available at https://github.com/CintiaG/MuPET-Flow/tree/master/example_data. The *S. pseudocapsicum* dataset is available http://flowrepository.org/id/FR-FCM-Z45W. *S. pastorianus* dataset was obtained directly from authors Gómez-Muñoz *et al*. (2021).

## ACKNOWLEDGMENTS

Flow cytometry data acquisition was performed at the IBPS Imaging Facility. The IBPS Imaging facility is supported by Region-Île-de-France, Sorbonne Université and CNRS.

