## Supplementary material for "MuPET-Flow: Multiple Ploidy Estimation Tool from Flow cytometry data"

**Table S1. Comparison of advantages and disadvantages for Cytoflow, flowPloidy and MuPET-Flow, tools used for measuring ploidy from flow cytometry data.**

| Cytoflow | flowPloidy | MuPET-Flow |
|---|---|---|
| **1. Upload data and configuration** | | |
| **Advantages**<br>• Graphical interface.<br>**Disadvantages**<br>• Requires manual configuration of experiment files. | **Disadvantages**<br>• Requires scripting for uploading the data and exploring the channels.<br>• The number of breaks for histograms needs to be known and specified. | **Advantages**<br>• Graphical interface.<br>• The available channels and number of breaks are automatically detected. |
| **2. Peaks' fluorescence intensity calculation** | | |
| **Advantages**<br>• It utilizes 1D mixture model, enabling the separation of the G0/G1 and G2 peaks and estimation of the mean intensity fluorescence more precisely.<br>**Disadvantages**<br>• It requires knowing the number of peaks *a priori*, and this same number is applied to all samples. | **Advantages**<br>• Automatic peaks and debris fitting using nonlinear regression.<br>• Offers the option to perform population gating and calculate the Coefficient of Variation. | **Advantages**<br>• The peaks are automatically calculated upon files upload.<br>• It uses a local maxima algorithm, which allows to detect and explore multiple peaks without previous information of the number of them.<br>**Disadvantages**<br>• Overlapping peaks can shift their fluorescence intensity. |
| **3. Histograms visualization and sample correction** | | |
| **Advantages**<br>• It allows to visualize individual as well as all samples simultaneously.<br>**Disadvantages**<br>• No curation option is available, although no errors were observed with our data. | **Advantages**<br>• Visualization of individual samples with a Shiny graphical interface.<br>**Disadvantages**<br>• Sometimes produces failed models that needed to be corrected by visualizing the samples individually and clicking on the peaks. | **Advantages**<br>• Visualization of individual and all samples enables quick detection and correction of noisy histograms if necessary.<br>**Disadvantages**<br>• Handling >36 samples may be challenging for visualization. |

| | | |
|---|---|---|
| **4. Histograms and peaks fluorescence export** | | |
| **Advantages**<br>• It allows exporting the histogram as an image and the peaks' fluorescence as a table. | **Disadvantages**<br>• The images and peaks' fluorescence export are via scripting. | **Advantages**<br>• Histograms can be exported as PNG or TIFF, and the peaks' fluorescence as a CSV table. |
| **5. Ploidy and genome size calculation** | | |
| **Disadvantages**<br>• The data needs to be exported, and ploidy must be calculated elsewhere (e.g., spreadsheet application or RStudio) | **Advantages**<br>• It can do automated genome size calculation in samples with internal standards.<br>**Disadvantages**<br>• If no internal standards were used, the calculated peaks fluorescence intensity needs to be exported, and ploidy or genome size calculated through additional operations. | **Advantages**<br>• It allows calculating ploidy or genome size within the app using a linear regression, and the results are reported in the same table of the peaks' fluorescence. |
| **6. General** | | |
| **Advantages**<br>• It can be utilized for various types of analyses.<br>**Disadvantages**<br>• Numerous manual configuration steps. | **Advantages**<br>• Powerful peaks and debris, which gives more precise results.<br>**Disadvantages**<br>• Heavily relies on scripting. | **Advantages**<br>• Simple to use and integrates all tools required for ploidy and genome size estimation.<br>• Multiple peaks detection.<br>**Disadvantages**<br>• No other type of analysis, such as gating or alternative plotting, can be performed. |

**Table S2. Conditions for developing and testing MuPET-Flow with the mock dataset.**

| **A. Hardware and environment.** | | | |
|---|---|---|---|
| **Computer Specifications** | **Software Environment** | **Execution Environment** | **Additional Information** |
| • Processor: Intel(R) Core(TM) i9-10900 CPU @ 2.80GHz.<br>• Memory: 32GB DIMM DDR4.<br>• Disk capacity: 4.5 TB. | • Operating System: Ubuntu 20.04.5 LTS.<br>• R version 4.2.2 Patched (2022-11-10 r83330). | • Code executed in RStudio 2022.02.3+492.<br>• Script executed in a single R session. | • No other CPU-intensive tasks were running during code execution.<br>• Parallel computing was not used in this analysis. |
| **B. Execution time.** | | | |
| **Process** | **User** | **System** | **Elapsed** |
| **Upload** | 1.23 | 0.05 | 1.28 |
| **Regression** | 0.02 | 0.00 | 0.02 |
| **Peaks** | 0.03 | 0.00 | 0.03 |
| **Plots** | 0.45 | 0.00 | 0.45 |
| **Total** | 1.72 | 0.05 | 1.78 |

**Table S3. MuPET-Flow results from the newly generated *Saccharomyces cerevisiae* dataset used to test the app for ploidy estimation.**

| Sample | Type | Channel | Smoothing | Window | Intensity G1 | Intensity G2 | Ploidy Mean | Ploidy Rounded |
|---|---|---|---|---|---|---|---|---|
| BY4742_rep1 | Standard | FL4-A | 0.04 | 10 | 100 | 173 | 1.00 | 1 |
| BY4742_rep2 | Standard | FL4-A | 0.1 | 50 | 110 | 190 | 1.00 | 1 |
| BY4742_rep3 | Standard | FL4-A | 0.1 | 50 | 122 | 203 | 1.00 | 1 |
| BY4743_rep1 | Standard | FL4-A | 0.1 | 50 | 192 | 343 | 2.00 | 2 |
| BY4743_rep2 | Standard | FL4-A | 0.1 | 50 | 212 | 379 | 2.00 | 2 |
| BY4743_rep3 | Standard | FL4-A | 0.1 | 50 | 239 | 415 | 2.00 | 2 |
| YPS128_3n_rep1 | Standard | FL4-A | 0.14 | 50 | 356 | 576 | 3.00 | 3 |
| YPS128_3n_rep2 | Standard | FL4-A | 0.1 | 50 | 297 | 507 | 3.00 | 3 |
| YPS128_3n_rep3 | Standard | FL4-A | 0.12 | 50 | 343 | 595 | 3.00 | 3 |
| YPS128_4n_rep1 | Standard | FL4-A | 0.14 | 50 | 486 | 808 | 4.00 | 4 |
| YPS128_4n_rep2 | Standard | FL4-A | 0.1 | 50 | 395 | 680 | 4.00 | 4 |
| YPS128_4n_rep3 | Standard | FL4-A | 0.14 | 50 | 463 | 803 | 4.00 | 4 |
| AVQ_rep1 | Test | FL4-A | 0.12 | 50 | 559 | 870 | 4.86 | 5 |
| AVQ_rep2 | Test | FL4-A | 0.1 | 50 | 471 | 793 | 4.97 | 5 |
| AVQ_rep3 | Test | FL4-A | 0.1 | 50 | 549 | 872 | 4.81 | 5 |
| CRE_rep1 | Test | FL4-A | 0.1 | 50 | 329 | 511 | 2.88 | 3 |
| CRE_rep2 | Test | FL4-A | 0.1 | 50 | 279 | 452 | 2.72 | 3 |
| CRE_rep3 | Test | FL4-A | 0.1 | 50 | 329 | 536 | 2.81 | 3 |

**Table S4. MuPET-Flow results from the *Saccharomyces pastorianus* dataset (Gómez-Muñoz *et al.*, 2021) used to test the app for ploidy estimation.**

| Sample | Type | Channel | Smoothing | Window | Intensity G1 | Intensity G2 | Genome Size Mean (Mb) | Genome Size Rounded (Mb) |
|---|---|---|---|---|---|---|---|---|
| *S. cerevisiae* CLA-1 | Standard | FL1-A | 0.1 | 50 | 93 | 176 | 12.10 | 12 |
| *S. eubayanus* FM1318 | Standard | FL1-A | 0.1 | 50 | 176 | 336 | 23.32 | 23 |
| *S. pastorianus* 1513 | Standard | FL1-A | 0.1 | 50 | 255 | 501 | 33.60 | 34 |
| *S. pastorianus* 1483 | Standard | FL1-A | 0.1 | 50 | 321 | 646 | 47.50 | 48 |
| *S. pastorianus* 790 | Test | FL1-A | 0.1 | 50 | 422 | 801 | 58.64 | 59 |

**Table S5. MuPET-Flow results from the S*olanum pseudocapsicum* dataset (Čertner *et al.*, 2022) used to test the app for samples with multiple peaks.**

| Sample | Type | Channel | Smoothing | Window | Intensity G1 | Intensity G2 | Ploidy Mean | Ploidy Rounded |
|---|---|---|---|---|---|---|---|---|
| *S. pseudocapsicum* mature-leaf | Standard | FL2-A | 0.1 | 50 | 390 | 821 | 2.00 | 2 |
| *S. pseudocapsicum* young-leaf | Standard | FL2-A | 0.12 | 50 | 387 | 803 | 2.00 | 2 |
| *S. pseudocapsicum* petal-blades | Test | FL2-A | 0.1 | 50 | 393 | 804 | 2.00 | 2 |
| *S. pseudocapsicum* root | Test | FL2-A | 0.1 | 50 | 396 | 828 | 2.04 | 2 |
| *S. pseudocapsicum* stem | Test | FL2-A | 0.1 | 50 | 408 | 829 | 2.06 | 2 |
| *S. pseudocapsicum* fruit-skin | Test | FL2-A | 0.1 | 50 | 1601 | 3203 | 7.68 | 8 |

**Table S6. Peaks' fluorescence intensity of all strains, replicates and peaks (Samples), and their ploidy obtained using different tools.**

| | Cytoflow | | flowPloidy | | MuPET-Flow | |
|---|---|---|---|---|---|---|
| **Samples** | **Intensity** | **Ploidy** | **Intensity** | **Ploidy** | **Intensity** | **Ploidy** |
| BY4742_rep1_G1 | 102.53 | 1.00 | 100.59 | 1.00 | 100.00 | 1.00 |
| BY4742_rep1_G2 | 170.06 | | 162.59 | | 173.00 | |
| BY4743_rep1_G1 | 201.25 | 2.00 | 190.66 | 2.00 | 192.00 | 2.00 |
| BY4743_rep1_G2 | 346.58 | | 337.03 | | 343.00 | |
| YPS128_3n_rep1_G1 | 350.04 | 3.00 | 350.75 | 3.00 | 356.00 | 3.00 |
| YPS128_3n_rep1_G2 | 567.83 | | 573.33 | | 576.00 | |
| YPS128_4n_rep1_G1 | 489.07 | 4.00 | 471.85 | 4.00 | 486.00 | 4.00 |
| YPS128_4n_rep1_G2 | 804.53 | | 811.33 | | 808.00 | |
| AVQ_rep1_G1 | 536.04 | 4.76 | 539.01 | 4.76 | 559.00 | 4.86 |
| AVQ_rep1_G2 | 870.88 | | 861.54 | | 870.00 | |
| CRE_rep1_G1 | 326.59 | 2.89 | 327.57 | 2.92 | 329.00 | 2.88 |
| CRE_rep1_G2 | 517.33 | | 515.91 | | 511.00 | |
| BY4742_rep2_G1 | 110.54 | 1.00 | 108.25 | 1.00 | 110.00 | 1.00 |
| BY4742_rep2_G2 | 192.10 | | 189.82 | | 190.00 | |
| BY4743_rep2_G1 | 220.27 | 2.00 | 213.95 | 2.00 | 212.00 | 2.00 |
| BY4743_rep2_G2 | 383.50 | | 379.49 | | 379.00 | |
| YPS128_3n_rep2_G1 | 295.51 | 3.00 | 293.53 | 3.00 | 297.00 | 3.00 |
| YPS128_3n_rep2_G2 | 509.71 | | 509.72 | | 507.00 | |
| YPS128_4n_rep2_G1 | 387.92 | 4.00 | 388.64 | 4.00 | 395.00 | 4.00 |
| YPS128_4n_rep2_G2 | 688.04 | | 683.60 | | 680.00 | |
| AVQ_rep2_G1 | 468.33 | 4.94 | 469.17 | 4.94 | 471.00 | 4.97 |
| AVQ_rep2_G2 | 796.21 | | 789.57 | | 793.00 | |
| CRE_rep2_G1 | 278.62 | 2.71 | 277.24 | 2.72 | 279.00 | 2.72 |
| CRE_rep2_G2 | 457.22 | | 454.58 | | 452.00 | |
| BY4742_rep3_G1 | 119.33 | 1.00 | 118.48 | 1.00 | 122.00 | 1.00 |
| BY4742_rep3_G2 | 202.22 | | 202.05 | | 203.00 | |
| BY4743_rep3_G1 | 243.45 | 2.00 | 243.83 | 2.00 | 239.00 | 2.00 |
| BY4743_rep3_G2 | 418.64 | | 417.49 | | 415.00 | |
| YPS128_3n_rep3_G1 | 335.35 | 3.00 | 334.97 | 3.00 | 343.00 | 3.00 |
| YPS128_3n_rep3_G2 | 588.95 | | 592.88 | | 595.00 | |
| YPS128_4n_rep3_G1 | 449.94 | 4.00 | 446.17 | 4.00 | 463.00 | 4.00 |
| YPS128_4n_rep3_G2 | 801.28 | | 793.92 | | 803.00 | |
| AVQ_rep3_G1 | 529.99 | 4.81 | 534.10 | 4.77 | 549.00 | 4.81 |
| AVQ_rep3_G2 | 895.58 | | 862.57 | | 872.00 | |
| CRE_rep3_G1 | 319.84 | 2.80 | 321.39 | 2.82 | 329.00 | 2.81 |
| CRE_rep3_G2 | 540.94 | | 540.04 | | 536.00 | |

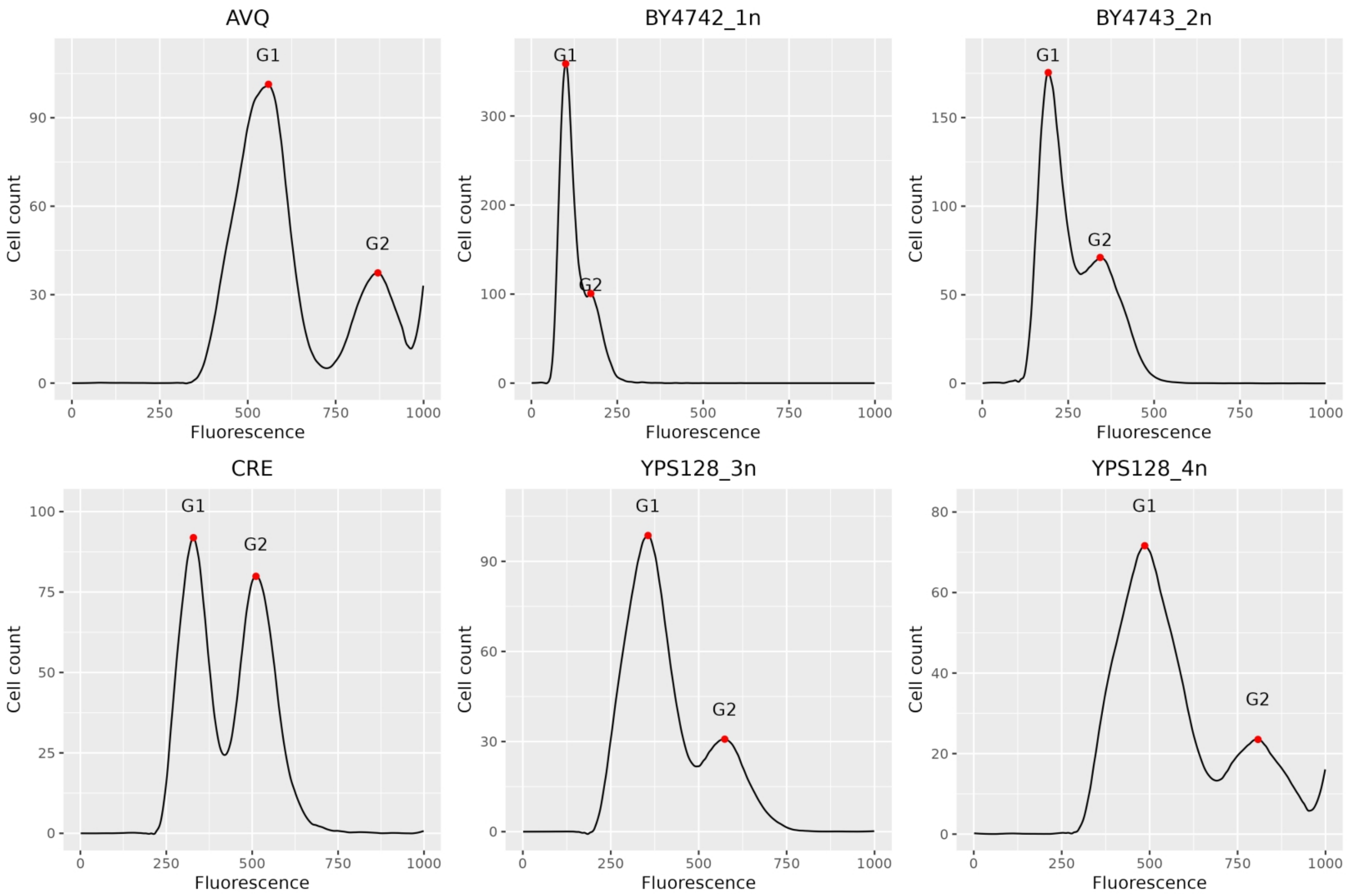


**Fig. S1. Fluorescence histograms of *Saccharomyces cerevisiae* strains of replicate 1, as obtained form MuPET-Flow (ordered alphabetically). Strains used as standards are shown with their ploidy as a suffix, and strains without a suffix are the tested strains. The ploidy inferred form the peaks' fluorescence of the test strains CRE and AVQ was of 3n and 5n, respectively.**
